# Pan-transcriptome-based Candidate Therapeutic Discovery for Idiopathic Pulmonary Fibrosis

**DOI:** 10.1101/824367

**Authors:** Yunguan Wang, Jaswanth K. Yella, Sudhir Ghandikota, Tejaswini C. Cherukuri, Harshavardhana H. Ediga, Satish K. Madala, Anil G. Jegga

## Abstract

**BACKGROUND:** Although the advent of two FDA-approved therapies for idiopathic pulmonary fibrosis (IPF) has energized the field, their effects are largely suppressive than pulmonary fibrosis remission- or reversion-inducing. Hence, the pursuit for newer IPF therapeutics continues. Recent studies show that joint analysis of systems biology level information with drug-disease connectivity are effective in discovery of biologically relevant candidate therapeutics.

**METHODS:** Publicly available gene expression signatures from IPF patients are used to query large scale perturbagen signature library to discover compounds that can potentially reverse dysregulated gene expression in IPF. Two methods are used to calculate IPF-compound connectivity: gene expression-based connectivity and feature-based connectivity. Identified compounds are further prioritized based on their shared mechanism(s) of action.

**RESULTS:** We identified 77 compounds as potential candidate therapeutics for IPF. Of these 39 compounds are either FDA-approved for other diseases or are currently in phase 2/3 clinical trials suggesting their repurposing potential for IPF. Among these compounds are multiple receptor kinase inhibitors (e.g., nintedanib, currently approved for IPF, and sunitinib), aurora kinase inhibitor (barasertib), EGFR inhibitors (erlotinib, gefitinib), calcium channel blocker (verapamil), phosphodiesterase inhibitors (roflumilast, sildenafil), PPAR agonists (pioglitazone), HDAC inhibitors (entinostat), and opioid receptor antagonists (nalbuphine). As a proof-of-concept, we performed *in vitro* validations with verapamil using lung fibroblasts from IPF and show its potential benefits in pulmonary fibrosis.

**CONCLUSIONS:** Since about half of the candidates discovered in this study are either FDA-approved or are currently in clinical trials for other diseases, rapid translation of these compounds as potential IPF therapeutics is feasible. Further, the generalizable, integrative connectivity analysis framework in this study can be readily adapted in early phase drug discovery for other common and rare diseases with transcriptomic profiles.

## Introduction

Idiopathic pulmonary fibrosis (IPF), a chronic and fatal fibrotic lung disease in people over 50 years old is estimated to affect 14 to 42.7 per 100,000 people (1). IPF is characterized by progressive subpleural and paraseptal fibrosis, heterogeneous honeycomb cysts (honeycombing), and clusters of fibroblasts and myofibroblasts (2). The median survival time of patients with IPF is 2.5 to 3.5 years, with 5-year survival rate around 20% (1). Currently, two small molecules - pirfenidone and nintedanib - are approved for IPF and are reported to slow down lung function decline caused by disease progression. However, drug induced side-effect profiles of these two drugs are formidable and their therapeutic effects are suppressive rather than pulmonary fibrosis remission or reversal (3, 4). The pursuit for relatively safer and efficacious therapies or combinatorials that arrest, or reverse fibrosis therefore continues. On the other hand, technological advances in experimental and computational biology resulted in rapidly expanding genomic and biomedical data, including transcriptomic profiles of disease and small molecules, disease or drug associated pathways and protein-protein interactions (5-7). Various approaches have been developed to facilitate *in silico* drug discovery via joint analysis of these data including the widely used connectivity mapping approach (8).

The concept of connectivity mapping between a drug and a disease is defined as the gene expression-based similarity calculated using a Kolmogorov-Smirnov statistic like algorithm (9, 10). It was first introduced as the ConnectivityMap (CMap) (8, 11) and succeeded by the LINCS L1000 project (CLUE platform) (8, 11), which currently contains gene expression profiles of ∼20,000 small molecule perturbagens analyzed in up to 72 cell lines. The connectivity mapping concept and application have led to the discovery of novel candidate compounds for disease, drug repurposing candidates, and novel drug mechanism of actions (12-16).

Recently, similar *in silico* drug discovery approaches for IPF have been reported wherein joint analysis of systems biology level information with drug-IPF connectivity are used to discover biologically relevant candidate therapeutics for IPF. For instance, Karatzas et al developed a scoring formula to evaluate drug-IPF connectivity obtained from multiple sources and identified several IPF candidate therapeutics (17). Interestingly, neither of the two approved IPF drugs (pirfenidone and nintedanib) was re-discovered by their approach (17). In another recent study using network-based approach and integrated KEGG network with connectivity analysis sunitinib, dabrafenib and nilotinib were identified as potential repurposing candidates for IPF (18).

Using a similar approach, namely, by examining connectivity between IPF gene signature and LINCS small molecules, we have previously reported 17-AAG (a known Hsp90 inhibitor) as a potential candidate therapeutic that inhibits fibroblast activation in a mouse model of pulmonary fibrosis (14). In another study, we screened connectivity of LINCS small molecules with cystic fibrosis (CF) and integrated with systems biology level information from CFTR to identify a candidate therapeutic for CF (13). These results suggest that disease-drug connectivity complemented with systems biology level information of drugs and disease could enable candidate therapeutic discovery. In the current study, we therefore calculated both gene-expression and enriched-pathway based connectivity between IPF and small molecules in an unsupervised manner and integrated these results with cheminformatics knowledge to prioritize candidate therapeutics for IPF. We identified 77 (out of ∼20,000 LINCS small molecules) candidate therapeutics for IPF. Significantly, among these 77 compounds was the approved drug for IPF (nintedanib), as well as several other compounds that are either currently being investigated or reported as a potential candidate therapeutics for IPF or investigated in clinical trial and reported to be ineffective (sunitinib, nilotinib, and sildenafil). *In vitro* and *in vivo* preclinical studies have reported beneficial effects of HDAC inhibitors (HDACIs) in preventing or reversing fibrogenesis (19, 20). Likewise, a previous study reported the beneficial effects of calcium channel blocking in bleomycin-induced pulmonary fibrosis (21). All these results suggest that the current approach has the potential to identify “true” candidate therapeutics. In the current study, we have selected verapamil, an FDA-approved calcium channel blocker, from our computational screening results for *in vitro* validation.

## Methods

### IPF studies/cohort selection

We used publicly available gene expression profiles from the Gene Expression Omnibus (GEO) (22) database for generating IPF gene expression signatures. Since gene expression profiles are known to be heterogeneous in different patients (23, 24), we selected 6 GEO datasets comparing primary healthy human lung tissues with primary IPF lung tissues for this study to potentially mitigate such heterogeneity (**Figure 1; Table 1**).

**Table 1:** Summary of 6 datasets comparing IPF lung tissue with healthy controls.

| GEO Data set | Sample description | Reference |
| --- | --- | --- |
| GSE10667 | 23 IPF samples and 16 controls | (38, 39) |
| GSE24206 | 17 IPF samples and 6 controls | (40) |
| GSE48149 | 13 IPF samples and 9 controls | (41) |
| GSE53845 | 40 IPF samples and 8 controls | (23) |
| GSE47460 | 131 IPF samples and 12 controls* | LGRC |
| GSE101286 | 7 IPF samples and 3 controls | (42) |

\* Only 12 control samples in the LGRC dataset with ‘normal’ clinical and pathological diagnosis were used as control.

**Figure 1.**
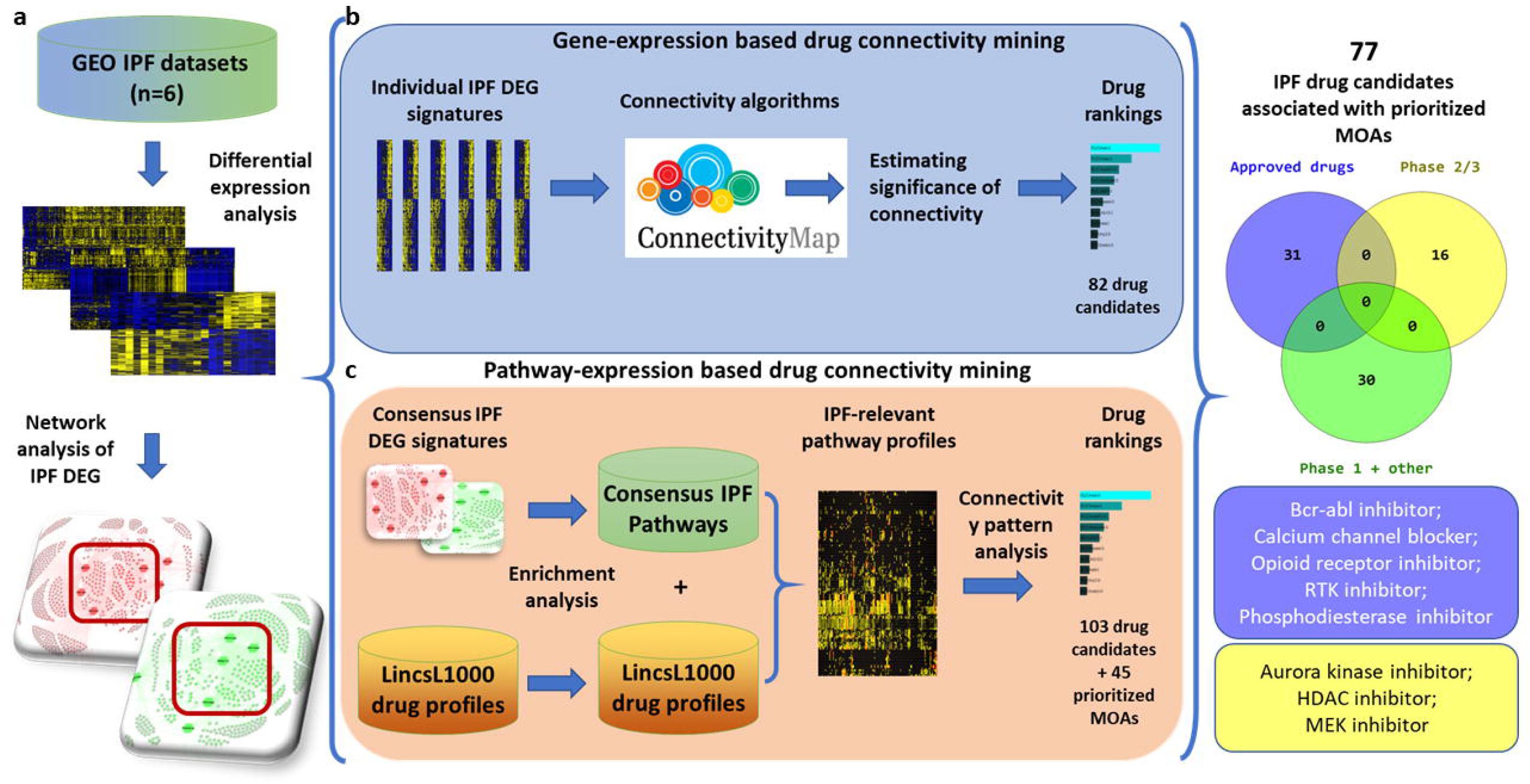
Overview of expression and annotation-based connectivity analysis. Workflow in this study could be summarized into three steps. a) Collection and differential analysis of human IPF gene expression datasets; b) Expression-based connectivity analysis through CLUE platform; c) annotation-based connectivity analysis examining similarity between pathways perturbed by compounds and those involved in IPF.

### Differential analysis of IPF gene expression profiles

Differential analysis was performed in R in using the package LIMMA (linear models for microarray data) (25). Genes with fold change ≥1.5 and adjusted p-value ≤0.05 were considered differentially expressed. Each dataset was analyzed separately.

### Known pulmonary fibrosis genes

We compiled 3278 “known” pulmonary fibrosis genes from literature and several data resources (**Supplementary Table 1**). This list contains human genes associated with “Pulmonary fibrosis”, “Idiopathic pulmonary fibrosis” and “Interstitial Lung Disease” from Open Targets platform (26), CTD (27), Phenopedia (28), and GeneCards (29) databases along with literature search (from PubMed).

### IPF-compound connectivity estimation and permutation analysis

To correct for multiple testing problem introduced by conducting connectivity analysis (**Figure 1**) in multiple datasets, we used permutation analysis to estimate the significance of connectivity. First, we constructed a matrix of connectivity score, denoted by *s*, between CLUE compound ***i*** and IPF dataset ***d*** in cell line ***j***. Next, positive, and negative connectivity to IPF were determined by thresholding connectivity score at 90 and -90, respectively:

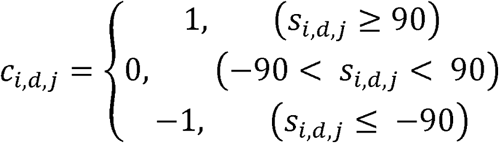

Overall connectivity, denoted by *o*, between each compound to IPF across all cell lines is summarized as the sum of individual connectivity across all datasets and all cell lines:

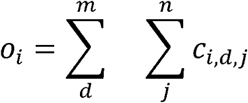

Permutation was performed by randomly shuffling rows of the connectivity matrix **C**, so that compound names were randomly assigned. Then, the permutated overall compound-to-IPF connectivity **O’** scores were calculated, and we recorded incidences where *o*_*i*_ ≤ *o*_*i*_, which indicates the observed compound to IPF connectivity is not larger than random connectivity:

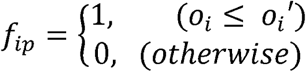

We repeated the permutation tests for 100,000 times and estimated significance as the frequency of **F** over all permutations. The significance cut-off was set at 0.05.

### Functional enrichment

Functional enrichment analysis was performed using pre-compiled gene annotation libraries from the ToppGene Suite (30). Enrichment p values were calculated using hypergeometric test in Python using the SciPy package.

### Annotation-based connectivity analysis

Annotation-based compound-IPF connectivity (**Figure 1**) was generated and evaluated as follows:

1. Identify enriched annotation terms in conserved IPF genes (genes that were differentially expressed in more than 4 IPF datasets) resulting in two vectors, ***I***_***up***_ and ***I***_***down***_;
2. Calculate enrichment score,***P***_***i,up***_ and ***P***_***i,down***_, using the 500 top and 500 bottom genes of each LINCS L1000 small molecule expression profile, denoted by **i**, in the up-regulated and down-regulated IPF pathways identified in step (1);
3. Calculate annotation-based connectivity score, defined as Pearson correlation between ***P***_***i,up***_ and ***P***_***down***_, and between ***P***_***i,down***_ and ***P***_***up***_;
4. To correct for false positives from multiple testing, permutation analysis were performed by swapping annotation terms in ***P***_***i,down***_ and ***P***_***i,up***_, followed up recalculation of annotation-based connectivity. 100,000 permutations were performed with significance threshold set to 0.05.

### Primary lung fibroblast cultures and RT-PCR

IPF lungs were collected in Dulbecco’s Modified Eagle Medium (DMEM) containing 10% FBS (Life Technologies, NY, USA) from the Interstitial Lung Disease Biorepository at the University of Michigan Medical School following the IRB regulations of the institute. Lung pieces were finely minced with sterile razor blades and incubated at 37°C for 30min in 5ml of DMEM containing collagenase (2mg/ml). Digested tissue was passed through a 100 µm filter, washed twice with DMEM medium, plated onto 100 mm tissue-culture plates, and incubated at 37°C, 5% CO_2_ to allow cells to adhere and migrate away from larger remaining tissue pieces. Adherent primary lung fibroblasts were harvested on Day 5 or 8 and lung-resident fibroblasts isolated with a negative selection using anti-CD45 beads as described earlier (JCI insight 2018). These fibroblasts were used for drug treatment studies up to passage four or less. After drug treatments, total RNA was extracted using RNAeasy Mini Kit (Qiagen Sciences, Valencia, CA, USA) and RT-PCR assays were performed. Relative quantities of mRNA for several genes were determined using SYBR Green PCR Master Mix (Applied Biosystems) and target gene transcripts in each sample were normalized to GAPDH and expressed as a relative increase or decrease compared with controls. As expected, we observed no changes in the copy number of GAPDH in IPF fibroblasts treated with verapamil compared to vehicle (0.0001 % DMSO). Also, we performed melt curve analysis to exclude primer sets producing non-specific PCR products. RT-PCR Primer sequences for genes, GAPDH (Fwd: AGCCACATCGCTCAGACAC; Rev: GCCCAATACGACCAAATCC), Col1α1 (Fwd: GGGATTCCCTGGACCTAAAG; Rev: GGAACACCTCGCTCTCCA), Col3α1 (Fwd: CTGGACCCCAGGGTCTTC; Rev: CATCTGATCCAGGGTTTCCA), Col5α1 (Fwd: CAGCCCGGAGAGAACAGA; Rev: GGTGCAGCTAGGTCATGTGAT), αSMA (Fwd: GCTTTCAGCTTCCCTGAACA; Rev: GGAGCTGCTTCACAGGATTC) and FN1 (Fwd: CTGGCCGAAAATACATTGTAAA; Rev: CCACAGTCGGGTCAGGAG).

## Result

### Differential expression analysis of IPF datasets

We analyzed 6 gene expression datasets comparing gene expression of IPF lung tissue with healthy controls (Table 1). Differential expression analysis was performed in each dataset using the R package LIMMA. Differentially expressed genes (DEG) were defined as genes with fold change above or equal to 1.5 and FDR-BH adjusted p-value less than or equal to 0.05. The number of DEGs ranged from 263 to 2385, and 4677 genes were unambiguously up-regulated, and 2210 genes were unambiguously down-regulated in at least one IPF dataset (**Figure 2a; Supplementary Table 1)**. Overall similarity between DEG gene lists was low, as reflected by the median Jaccard index between gene lists (0.077). While the lack of concordance between datasets suggests disease heterogeneity in IPF, it also provides the rationale for meta-analysis using multiple datasets to extract high confidence drug candidates for IPF. Despite the overall heterogeneity in DEG among different IPF datasets, there were also a considerable number of genes that were consistently dysregulated in 4 or more IPF datasets. We call these genes as “conserved” IPF genes (197 up-regulated genes and 84 down-regulated genes). Among these conserved DEGs, 179 genes (121 upregulated and 58 downregulated) were known previously to be involved in pulmonary fibrosis (**Figure 2b)**. Functional enrichment analysis of conserved IPF genes showed that biological processes involved in extracellular matrix formation, inflammation responses and cell migration were up-regulated while processes involved in normal lung processes such as angiogenesis and alveolar functions were down-regulated.

**Figure 2.**
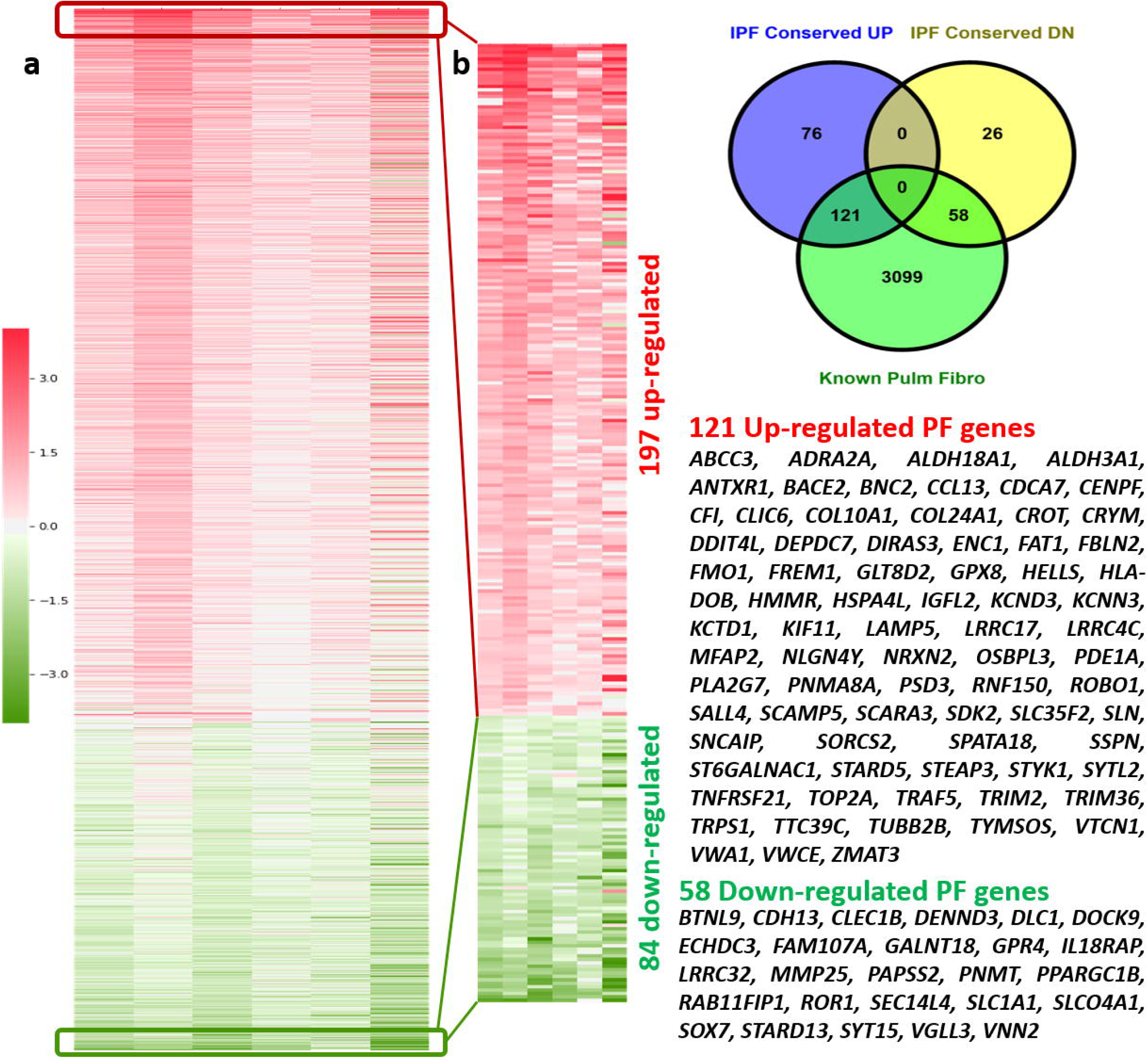
Heat map view of differentially expressed genes in 6 IPF datasets. Expression of 2206 genes unambiguously differentially expressed in at least two IPF dataset are shown. Genes are represented in rows and patient samples in columns. Cells in heat map were sorted discerningly based on median log fold change in 6 datasets and on number of datasets they were differentially expressed in. b) Genes that were upregulated or downregulated in at least 4 datasets. Intersection with all known pulmonary fibrosis genes are shown in the Venn diagram.

### Expression based connectivity analysis and permutation analysis

We used the NIH Library of Integrated Network-Based Cellular Signatures (LINCS) library as the compound search space for IPF candidate therapeutics. The LINCs Touchstone dataset contains a total of approximately 8,400 perturbagens, including more than 2000 small molecules that have produced gene signatures generated from testing on a panel of nine cell lines. These cell lines include A375, A549, HEPG2, HCC515, HA1E, HT29, MCF7, PC3, and VCAP. The LINCS library contains expression profiles of ∼20,000 small molecules assayed in various cell lines. To identify potential IPF candidate therapeutics, we adopted the connectivity mapping method which assumes small molecules with gene expression profiles negatively correlated with that of a disease are likely to be therapeutic for the disease. We first queried the Connectivity Map web platform (CLUE.io) for compounds with a reversed gene expression profile comparing to IPF. From each dataset, a gene signature containing up to 150 most up-regulated and down-regulated genes was extracted and used to query the CLUE platform (**Supplementary Table 2)**. Using the hits from CLUE results, we applied a “greedy” approach to capture the highest number of compounds connected to IPF by selecting compounds with at least 90 connectivity score in any one of the cell lines. This approach returned 1000+ compounds that relate to IPF at least once. However, this approach results in several compounds that are connected to IPF in both the directions, i.e., potentially inducing, and reversing IPF gene expression profiles at the same time (**Figure 3a)**. This suggests that the gene expression perturbation due to technical variation is present in our data and is reflected in the form of these low frequency compounds in CLUE analysis. Based on the assumption that IPF-related gene expression patterns are consistently present in our selected 6 IPF datasets, we performed permutation analysis to estimate the significance of IPF-disease connectivity and filter out potential false positives. With a 0.05 significance cut-off, we identified 82 compounds that were significantly connected with IPF (**Figure 3b)**. These compounds were associated with 63 different known drug mechanisms of action.

**Figure 3.**
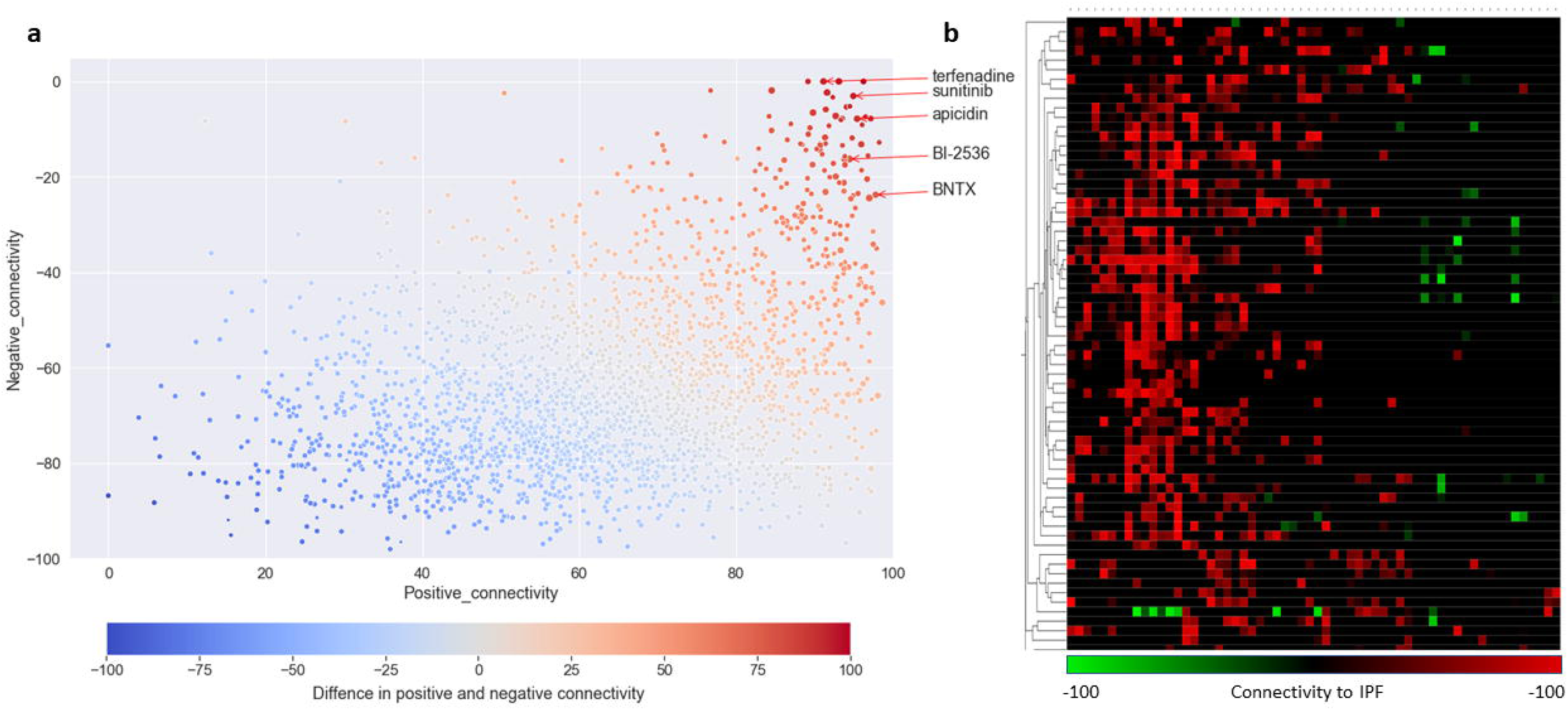
Gene expression-based connectivity score of CLUE compounds. Scatter plot of positive connectivity score against negative connectivity score between each CLUE compound and each IPF dataset across all cell lines. Coordinates of each point were determined by the average of highest or lowest 6 connectivity scores among all 54 values across 6 IPF datasets and 9 cell lines in CLUE. b) Heat map view of connectivity score of 82 compound that were significantly connected to IPF based on permutation analysis.

### Functional enrichment-based connectivity analysis

A major limitation of using the Touchstone Library-based CLUE platform to screen for drug candidates is that it only covers 2,836 out of ∼20,000 small molecules available in the full L1000 database. To overcome this, we searched for IPF candidate therapeutics among the remaining ∼17,000 LINCS small molecules. To do this, we used a functional enrichment based metric to evaluate drug-IPF connectivity of these LINCS small molecules, wherein gene expression data from both the compound and the IPF data sets were transformed into enrichment p-values using hypergeometric tests against gene functional annotations (Gene Ontology Biological Process, Mouse Phenotypes and KEGG pathways). To minimize the noise introduced by biological processes not relevant to IPF, we only considered functional annotations enriched in the conserved IPF gene sets, and thus, each compound enrichment profile was limited to these pathways. Connectivity between a compound and IPF was defined as the cosine similarity between the p-values of pathways enriched in compound up-regulated genes and those enriched in IPF down-regulated genes, and vice versa. Next, we used permutation analysis as discussed in the previous sections to estimate the significance of the feature-based compound-IPF connectivity and identified 345 compounds that perturbed IPF-related pathways in an overall opposite manner compared to IPF. To find groups of functionally related therapeutic candidates that could act on IPF perturbed biological processes, we performed clustering analysis on the compound enrichment profiles and prioritized four clusters of compounds with high connectivity to IPF. The compounds in three clusters selectively down regulated pathways such as ‘cell adhesion’, ‘collagen metabolic process’ and ‘regulation of programmed cell death’, which were all upregulated in IPF. On the other hand, compounds in the cluster that showed up-regulation of ‘blood vessel morphogenesis’ and ‘angiogenesis’ were down regulated in IPF. The approved IPF drug, nintedanib was in this cluster of compounds. Combining compounds from these four clusters, we identified 103 candidates as IPF therapeutic candidates from annotation- or feature-based connectivity analysis (**Figure 4)**. These compounds include EGFR inhibitor gefitinib, PDGFR and VEGFR inhibitor dovitinib, and KIT inhibitor sunitinib.

**Figure 4.**
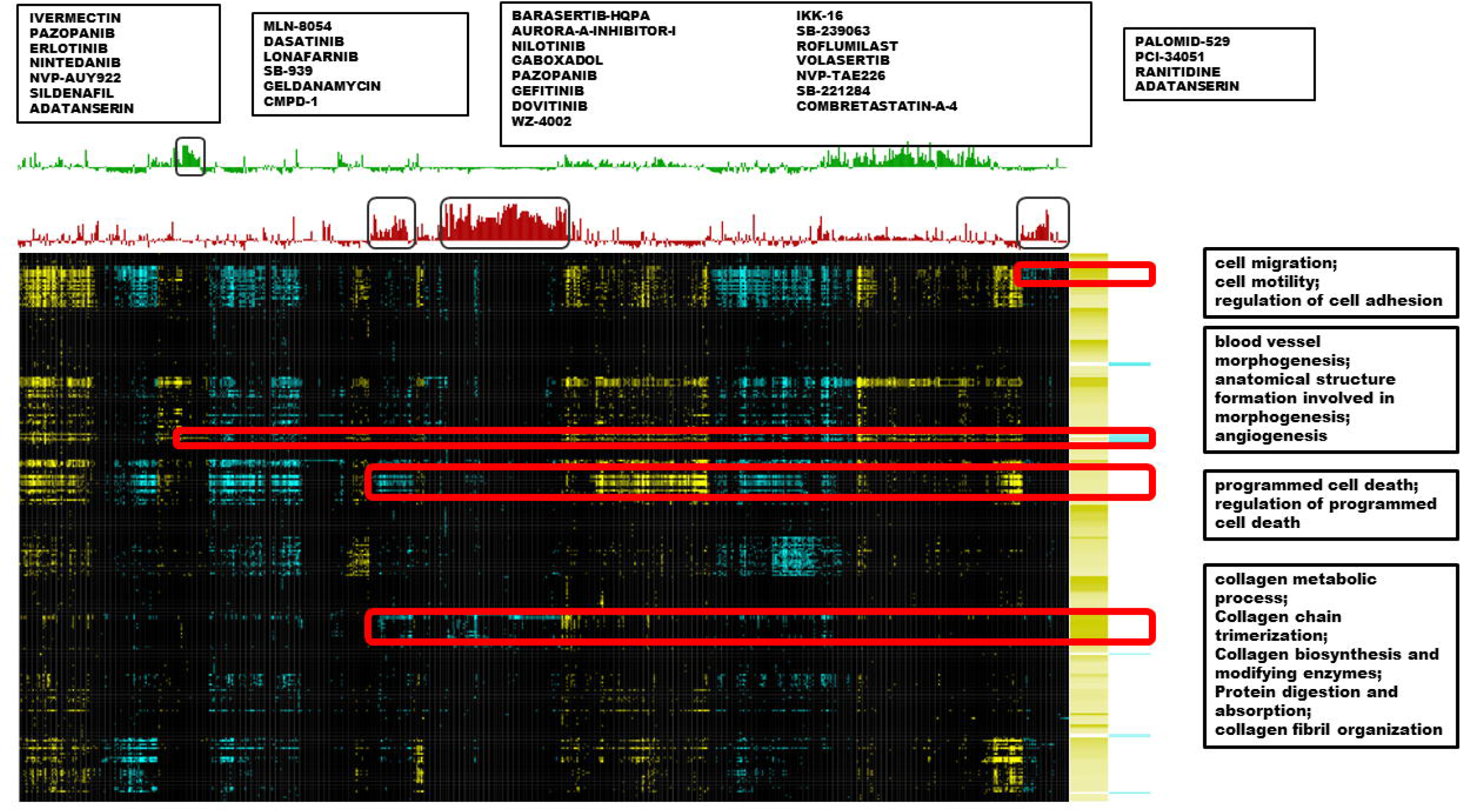
Enriched pathway heat map view of 345 drugs significantly connected to IPF in annotation-based connectivity analysis. Enrichment terms in categories including KEGG pathways, Wiki pathways, REACTOME, Mouse phenotype and Gene Ontology: Biological process in either consistently up-regulated or down-regulated IPF genes are arranged in rows. LINCS drug profiles were arranged in columns. Blue indicates annotation terms are enriched in down-regulated genes by drug or IPF, and yellow indicates annotation terms are enriched in up-regulated genes by drug or IPF. K-means clustering was used to identify compound modules.

### Prioritization of IPF candidate therapeutics based on shared mechanism of action

In the annotation-based connectivity analysis, we observed that most of the discovered compounds shared drug mechanisms of action. For instance, EGFR inhibition, PDGFR inhibition and VEGFR inhibition were shared across multiple compounds suggesting potential relevance of these MoAs to IPF. This also suggests a likelihood of higher therapeutic potential of multiple compounds with shared or similar MoAs. Leveraging the known MoA information of the discovered compounds, we further prioritized compounds belonging to MoA that were prioritized by annotation-based connectivity analysis. After removing glucocorticoid receptor agonists and immunosuppressants from the list because of their known detrimental effects in IPF, we identified 48 MoAs meeting these criteria (**Supplementary Table 3**). Based on these MoAs, we selected 77 compounds as our final pre-clinical candidates for IPF (**Supplementary Table 4)**. Among these, 39 were FDA-approved drugs or phase-II/III compounds suggesting their repositioning potential for IPF (**Table 2)**. These drugs include Bcr-Abl kinase inhibitors, EGFR inhibitors, opioid receptor inhibitors, receptor tyrosine kinase (RTK) inhibitors and aurora kinase inhibitors. Notably, the approved IPF drug nintedanib was also among this list. This is important because nintedanib was not included in the CLUE database and would have been missed if annotation-based connectivity were not examined. A closer look at the pharmacological targets of these candidates revealed that many of these targets such as PDGFRA, EGFR, FGFR4, FYN and KDR were differentially expressed in IPF. Similarly, CACNA1G, SLC29A4, CACNB3, SLC6A4 – all targets of verapamil, a known calcium channel blocker, were differentially expressed in IPF. Among these targets, KDR, FGFR4 and PDGFRA are associated with nintedanib and other multi-targeted RTK inhibitors such as dovitinib, pazopanib and sunitinib (**Figure 5)**. These genes are involved in VEGF and PI3K/AKT signaling and VEGFR2 mediated cell proliferation, suggesting a role for multi-targeted RTK inhibitors in controlling IPF through VEGF signaling inhibition.

**Table 2:** List of 39 potential repurposing candidates for IPF.

| Compound | IPF-related MOA | Indication | Phase |
| --- | --- | --- | --- |
| febuxostat | xanthine oxidase inhibitor | hyperuricemia | Launched |
| nortriptyline | tricyclic antidepressant | depression | Launched |
| amsacrine | topoisomerase inhibitor | cancer | Launched |
| irinotecan | topoisomerase inhibitor | cancer | Launched |
| camptothecin | topoisomerase inhibitor | cancer | Phase 3 |
| pioglitazone | ppar receptor agonist, insulin sensitizer | diabetes mellitus | Launched |
| roflumilast | phosphodiesterase inhibitor | COPD | Launched |
| sildenafil | phosphodiesterase inhibitor | Erectile dysfunction and pulmonary hypertension | Launched |
| nalbuphine | opioid receptor antagonist | pain relief | Launched |
| everolimus | mTOR inhibitor | cancer | Launched |
| sunitinib | mRTK inhibitor | cancer | Launched |
| nintedanib | mRTK inhibitor | IPF | Launched |
| dovitinib | mRTK inhibitor | cancer | Phase 3 |
| pazopanib | mRTK inhibitor | cancer | Launched |
| selegiline | monoamine oxidase inhibitor | Parkinson's Disease | Launched |
| selumetinib | mek inhibitor | cancer | Phase 3 |
| curcumin | lipoxygenase inhibitor, histone acetyltransferase inhibitor, cyclooxygenase inhibitor |  | Launched |
| tomelukast | leukotriene receptor antagonist | asthma | Phase 3 |
| dasatinib | mRTK inhibitor | cancer | Launched |
| lafutidine | histamine receptor antagonist | duodenal ulcer disease, peptic ulcer disease (PUD) | Launched |
| ranitidine | histamine receptor antagonist | heartburn | Launched |
| amodiaquine | histamine receptor agonist | malaria | Launched |
| entinostat | hdac inhibitor | cancer | Phase 3 |
| remacemide | glutamate receptor antagonist | epilepsy and neurodegenerative diseases | Phase 3 |
| riluzole | glutamate inhibitor | amyotrophic lateral sclerosis (ALS) | Launched |
| erlotinib | egfr inhibitor | cancer | Launched |
| gefitinib | egfr inhibitor | cancer | Launched |
| sulpiride | dopamine receptor antagonist | schizophrenia | Launched |
| mycophenolic acid | dehydrogenase inhibitor, inositol monophosphatase inhibitor | organ rejection | Launched |
| ketorolac | cyclooxygenase inhibitor | NSAID | Launched |
| mestinon | cholinesterase inhibitor | myasthenia gravis | Launched |
| fipronil | chloride channel blocker | insecticide | Launched |
| verapamil | calcium channel blocker | hypertension | Launched |
| gaboxadol | benzodiazepine receptor agonist | insomnia | Phase 3 |
| ivermectin | benzodiazepine receptor agonist | gastrointestinal parasites | Launched |
| nilotinib | bcr-abl kinase inhibitor | cancer | Launched |
| barasertib-HQPA | aurora kinase inhibitor | cancer | Phase 2/Phase 3 |
| regadenoson | adenosine receptor agonist | myocardial perfusion imaging (MPI) | Launched |
| bucladesine | adenosine receptor agonist | skin ulcer | Launched |

**Figure 5.**
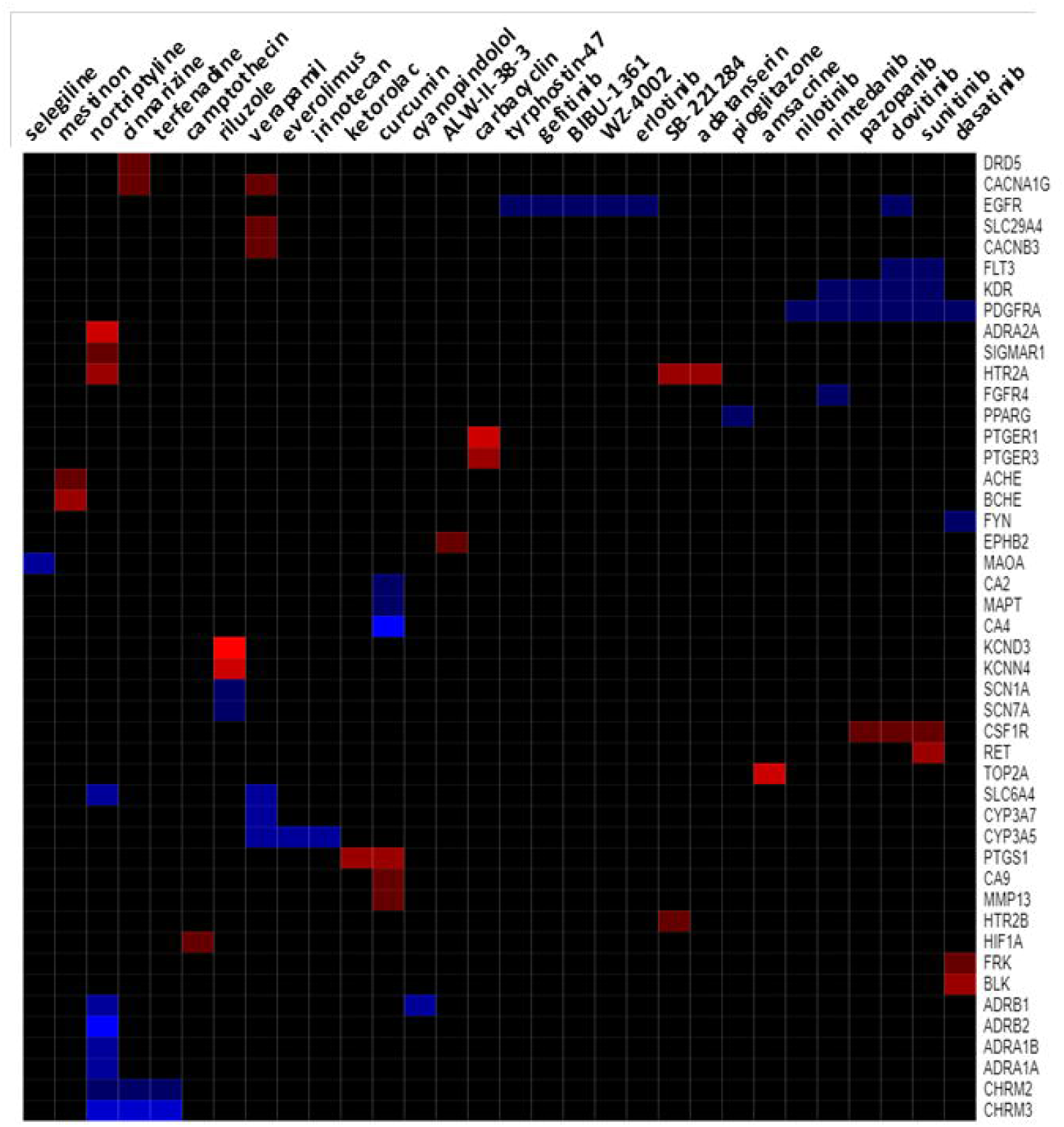
Heat map view of candidate compound targets that are differentially expressed in IPF. Log fold changes of 48 target genes of 30 compounds in IPF are shown. Only compounds with targets differentially expressed in at least two IPF datasets are included. Differentially expressed drug targets in IPF are in rows and discovered IPF candidate compounds are in columns. Rows and columns are ordered using 2D hierarchical clustering.

### Anti-fibrotic potential of verapamil

Calcium channel blockers are commonly used in clinical practice and reported to be well-tolerated. Therefore, in the current study, as a proof-of-concept, we have selected verapamil, a known calcium channel blocker and anti-hypertension drug, from our computational screening results for *in vitro* validation. IPF lung fibroblasts were treated with either vehicle (0.001% DMSO) or verapamil (Figure 6). We observed a significant reduction in the expression of fibroproliferative genes (*FN1, COL1A1, COL3A1*, and *COL5A1*), and αSMA with verapamil treatment for 16 hrs. compared to vehicle treated IPF fibroblasts (**Figure 6**). These findings support the premise that candidate small molecules identified using *in silico* screening methods are potentially effective in inhibiting fibroblast activation and may serve as potential drug candidates for further validation using *in vitro* and *in vivo* pre-clinical IPF models. Testing of additional candidates however is warranted.

**Figure 6.**
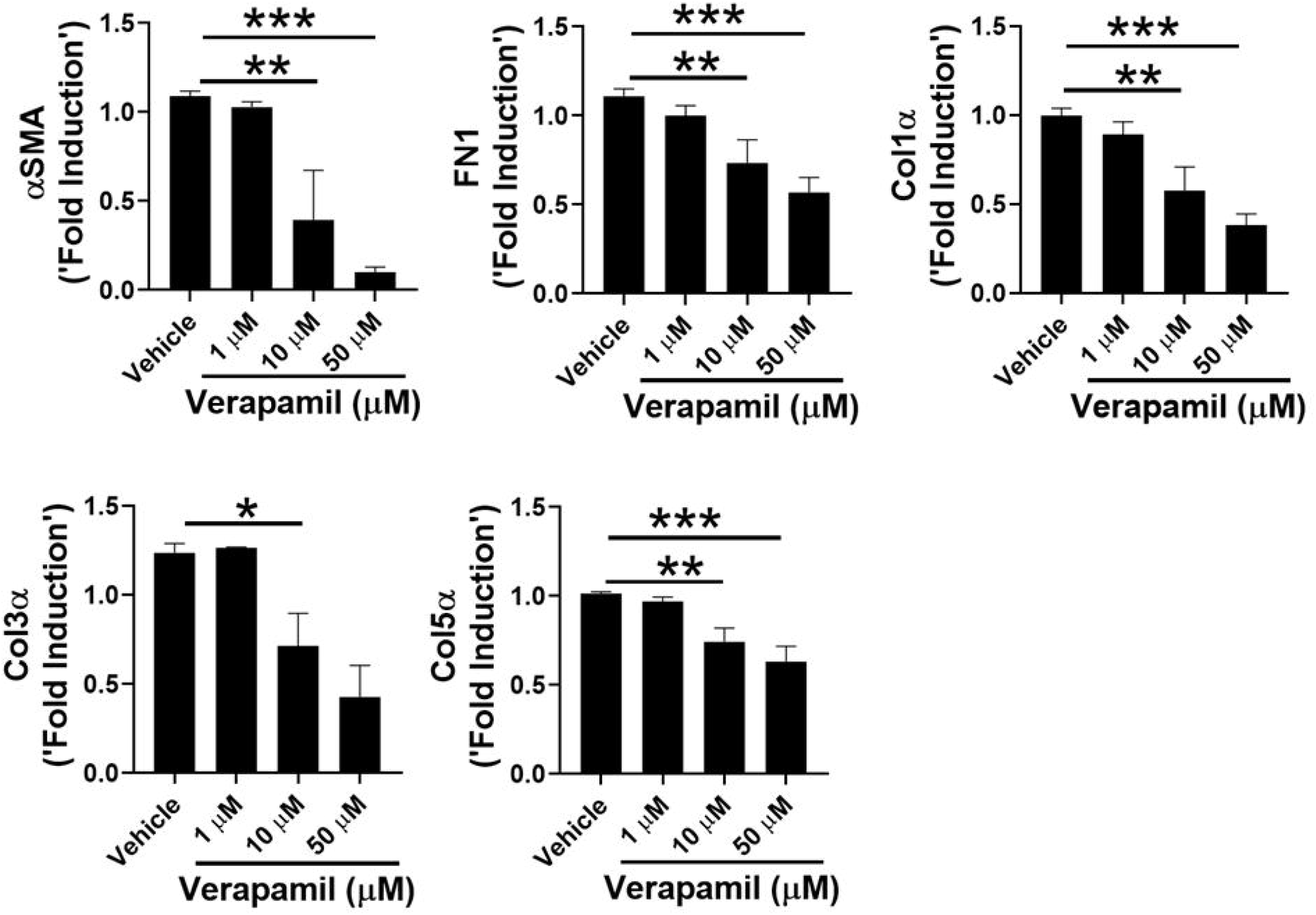
Verapamil treatment attenuates pro-fibrotic gene expression. IPF fibroblasts were treated with either vehicle (DMSO 0.001%) or verapamil (1, 10 and 50 µM) for 16 hrs. Total RNA was analyzed for the expression of αSMA and ECM genes (FN1, Col1α, Col3α and Col5α) using RT-PCR. *P<0.05; **P<0.005 ***P<0.0005; ****P<0.00005.

## Discussion

In this study, we developed a multiplexed, generalizable approach to discover novel therapeutics by integrating disease-driven and perturbagen-driven gene expression profiles, disease-associated biological pathways, and cheminformatics of perturbagen. Compound-IPF connectivity was examined from different dimensions including transcriptome, functional enrichment profiles, and drug mechanisms of actions. With this approach, we not only identified approved IPF therapeutic drugs (nintedanib) but also identified additional FDA-approved drugs that share similar MOA as IPF candidate therapeutics. Notably, these drugs were not discoverable using conventional transcriptome-based connectivity analysis alone. Further, approved drugs or investigational compounds associated with MoAs such as Bcr-abl inhibitor and Aurora kinase inhibitor, were also among our candidate list, which could provide insights into novel intervention strategies against IPF.

Transcriptome based connectivity mapping was first introduced more than a decade ago and has been applied to facilitate drug discovery for various diseases, including IPF. In a recent study, Karatzas et al proposed nine drugs as IPF therapeutics based on their connectivity with expression profiles derived from IPF datasets. However, they were not able to re-discover approved IPF drugs although these drugs were included in the LINCS L1000 library that was queried (17). Likewise, our gene expression-based connectivity approach through CLUE prioritized 82 small molecules as IPF therapeutics candidates, but neither of the FDA approved IPF drugs (pirfenidone or nintedanib) was in the list. A closer look at the connectivity scores revealed that while pirfenidone, one of the two approved IPF therapeutics, had no connectivity to IPF in any of the six queried datasets, pirfenidone was connected to verapamil (transcriptionally similar – based on connections/internal connectivities in Clue.io platform). Our pre-clinical in vitro validations with verapamil using human IPF lung fibroblast revealed therapeutic benefit of verapamil in IPF. A previous study reported the beneficial effects of calcium channel blocking in bleomycin-induced pulmonary fibrosis (21). Based on these preliminary results and data, we hypothesize that verapamil and pirfenidone combination could have potential therapeutic benefit in IPF patients. Additional in vitro and in vivo combined (pirfenidone plus verapamil) vs monotherapy (verapamil) are however required to test and confirm this hypothesis. Additionally, mining electronic health records and side-effects data (FAERS FDA’s adverse events reporting system) for testing whether patients under both therapy have a better response are part of our related ongoing studies (31). Nintedanib, the second approved therapeutic for IPF, was not included in CLUE and therefore we were unable to assess its connectivity to IPF using Clue.io platform.

We also examined the connectivity between IPF and the ∼17,000 compounds not covered in the Clue.io platform. The gene expression profiles associated with these ∼17K small molecules are from distinct selections of cell lines. Therefore, direct connectivity mapping analysis may be susceptible to biological variation introduced by different cell lines. In addition, it has been shown that integration of prior knowledge, particularly in the form of gene sets information in biological pathways, improves the accuracy of drug activity predictions (32). We evaluated drug-IPF connectivity through enriched pathways directly related to IPF under the assumption that pathways perturbed by drugs are more stable across different host cell conditions compared to individual genes. Annotation-based connectivity analysis lead to discovery of additional 14 small molecules that were not included in the CLUE platform. These include aurora kinase inhibitor barasertib-HQPA and phosphodiesterase inhibitor roflumilast. Notably, nintedanib was also among the 14 additional small molecules, and the enriched pathways that contributed to the connection to IPF were related to fibroblast proliferation, ECM production and cell migration, which is in consistent with implicated MoA of nintedanib against IPF *in vitro* (33).

Among the 77 prioritized candidates, 31 are FDA-approved drugs and are associated with different MoAs. These MoAs include RTK inhibition, which is the known MoA for the approved IPF drug nintedanib. Other compounds with this MoA in our discovered candidate compounds include pazopanib and sunitinib. Sunitinib is approved for treatment of renal cell carcinoma and gastrointestinal stromal tumor, and it has also been shown to be efficacious in inhibiting established pulmonary fibrosis in the bleomycin-induced mouse model (34). Additionally, MoAs involved with these 31 drugs also included those associated with compounds that are currently investigated or are in clinical trial for IPF drugs, such as src-kinase inhibitor and mTOR inhibitor (35). The MoAs associated with the remaining IPF repositioning candidates included aurora kinase inhibitor, EGFR inhibitor, calcium channel blocker, phosphodiesterase inhibitor, PPAR agonist, Bcr-Abl kinase inhibitors and opioid receptor antagonist. HDACIs have been reported to improve resolution of pulmonary fibrosis in mice (19, 20). Calcium channel blockers like verapamil are commonly used in clinical practice, reported to be well-tolerated, and are relatively inexpensive. Further, a previous study using bleomycin model of pulmonary fibrosis has reported its benefits (21).

While the computational drug discovery approaches, including the current one, are powerful approaches for pre-clinical therapeutic discovery and MoA-based hypotheses, they albeit suffer with certain inherent limitations. For example, sildenafil, one of the 77 compounds that we have discovered as candidate therapeutics for IPF, has already been investigated in combination with nintedanib in clinical trials (36, 37) and was reported to have no significant benefit when compared to patients on nintedanib alone. Nevertheless, discovering a compound (sildenafil) that is tested in a clinical trial for IPF demonstrates the preclinical discovery power of our approach for candidate therapeutics. Second, the current approach is predominantly monotherapy-centric and does not consider potential drug-drug interactions. For example, the current approach cannot deduce which of the 77 compounds can be a potential combination compound with nintedanib or pirfenidone. Advanced knowledge-mining (e.g., known drug-drug interactions) and machine learning based approaches can be potentially explored to address this problem. Third, ours, and other computational approaches do not consider the potential off-target effects leading to adverse events. For example, as a previous study too reported (21), off-target effects of calcium channel blockers on other cell types cannot be ruled out especially because the lung is composed of more than 50 different cell types with several of these expressing voltage-dependent calcium channels.

In conclusion, we have developed a generalizable, integrative connectivity analysis combining information from transcriptomic profiles, disease systems biology and drug cheminformatics for *in silico* IPF drug discovery. Application of our approach earlier in the IPF drug discovery pipeline may help to avert late stage clinical trial failures. As almost half the candidates we have discovered in this study are FDA-approved or are currently in clinical trials for several diseases, rapid translation of these compounds is feasible. Finally, we have suggested novel drug mechanisms that could shed new insights in the search for better IPF drugs.

## DECLARATIONS

### Ethics approval and consent to participate

Not applicable.

### Consent for publication

Not applicable.

### Availability of data and materials

All data generated or analyzed during this study are included in this published article and its supplementary information files.

### Author contributions

Y.W. and A.J. conceived this study. Y.W., J.Y. and S.G. collected and analyzed data. Y.W. and A.J. interpreted results from data. T.C., H.E., and S.M. conducted experimental validations. Y.W., S.M. and A.J. edited this manuscript. Y.W. wrote the first draft.

## Acknowledgements

Not applicable.

